# Chi-miR-3880 mediates the regulatory role of interferon gamma in goat mammary gland

**DOI:** 10.1101/2022.10.10.511576

**Authors:** Yue Zhang, Jidan Liu, Guanglin Niu, Qiong Wu, Fangjun Cao, Binyun Cao

**Affiliations:** College of Animal Science and Technology, Northwest A&F University, 712100 Yangling, Shaanxi, China; Longmen Animal Disease prevention and Control Center, 516800 Huizhou, Guangdong, China; Department of Oncology Pathology, Karolinska Institutet, 17164 Stockholm, Sweden; School of Life Sciences, Technical University of Munich, 85354 Freising, Germany; Medical College, Qinghai University, 810001 Xining, Qinghai, China

**Keywords:** interferon gamma, chi-miR-3880, interferon stimulated genes, goat lactation

## Abstract

A healthy mammary gland is a necessity for milk production of dairy goats. The role of chi-miR-3880 in goat lactation is illustrated in our previous study. Among the differentially expressed genes regulated by chi-miR-3880, one seventh are interferon stimulated genes, including *MX1*, *MX2*, *IFIT3*, *IFI44L*, and *DDX58*. As the inflammatory cytokine interferon gamma (IFNγ) has been identified as a potential marker of caseous lymphadenitis in lactating sheep, the interaction between IFNγ and immune-related microRNAs is explored. In this study, it was found that chi-miR-3880 could be one of the microRNAs downregulated by IFNγ in goat mammary epithelial cells (GMECs). The regulation among IFNγ, chi-miR-3880, and *DDX58*, as well as their roles in mammary gland were investigated. The study illustrated that IFNγ/chi-miR-3880/*DDX58* axis modulates GMEC proliferation and lipid formation through PI3K/AKT/mTOR pathway, and regulates apoptosis through Caspase-3 and Bcl-2/Bax. The role of the axis in involution was reflected by the expression of p53 and NF-κB. In conclusion, IFNγ/chi-miR-3880/*DDX58* axis plays an important part in lactation.

## Introduction

The health of the mammary glands is one of the recognized factors for milk production in dairy goats. MicroRNAs are a family of short non-coding RNAs, ~22 nucleotides in length, that mediate target gene silencing by binding to the target sites of mRNAs in the 3’ untranslated region (UTR) [1, 2]. Indirectly, microRNAs are involved in the regulation of the cellular processes, which could be essential in animal development [2]. In previous studies, it is found that microRNAs play important roles in goat reproductive characteristics, such as fertility and lactation [3–8]. Chi-miR-3880 is one of the microRNAs upregulated during goat peak lactation period compared to colostrum period that facilitates proliferation and lipid accumulation of goat mammary epithelial cells (GMECs), promotes mammary gland development, and extends the length of lactation period [7]. In this study, the differentially expressed genes induced by chi-miR-3880 [7] were analyzed. Strikingly, it was found that ~14% of chi-miR-3880 regulated differentially expressed genes are interferon stimulated genes (ISGs).

As one of the downstream genes of chi-miR-3880 [7], *DExD/H-Box Helicase 58 (DDX58*), also known as retinoic-acid-inducible gene I (*RIG-I*), is a cytoplasmic viral RNA receptor that recognize viral double-stranded RNAs to elicit antiviral IFN responses [9, 10]. An interferon (IFN) response is described to be related to goat reproduction [11] and protect mammals from viral infection [12]. During infection, the production of IFNs could be lifted, which induces the increased expression of related genes, typically ISGs [13]. IFN gamma (IFNγ), a type II IFN, is one of the secreted cytokines with an important regulatory role in immunity and inflammation, which has been identified as a potential marker of caseous lymphadenitis in lactating sheep [14]. In this study, the regulation among IFNγ, chi-miR-3880, and *DDX58*, as well as their roles in mammary gland were investigated.

## Materials and Methods

### Animal Ethics

All procedures conformed to institutional and national guidelines, and were approved by the Experimental Animal Management Committee of Northwest A&F University (ethic code: #0726/2018).

### Cell Cuture

Goat mammary epithelial cells (GMECs) were isolated from mammary gland tissue of Guanzhong dairy goats during lactation period. Specifically, fresh mammary gland tissue was washed off milk with phosphate buffer saline and surface-sterilized with 75% ethanol. The tissue was then cut into 1 mm^3^ cubes to seed in 35 mm cell culture dishes. GMECs that spread out around the tissue cubes were purified by trypsinization. GMECs were cultured in DMEM/F12 medium (Hyclone, Waltham, MA, USA) with 10% fetal bovine serum (Gibco, United States) and penicillin/streptomycin, and incubated in 5% CO_2_ at 37°C in a humid atmosphere. The miRNA and siRNA were synthesized at GenePharma Corporation (Shanghai, China). The sequence of chi-miR-3880 is 5’-GGUCCCGCCGCCGCCGCC-3’, and that of si*DDX58* is 5’-GGUACAAAGUUUCAGGCAU-3’. The negative control (NC) was designed and synthesized by GenePharma Corporation (Shanghai, China): 5’-UUCUCCGAACGUGUCACGU-3’. Small RNAs were transfected into GMECs at a final concentration of 50 nM using Lipofectamine RNAiMAX reagent (Invitrogen, USA) when the cells reach ~80% confluence.

### RNA sequencing data analysis

The downstream differentially expressed genes regulated by chi-miR-3880 [7] were enriched to GO terms and KEGG pathways. The GO enrichment was performed at http://geneontology.org/, and the KEGG pathway enrichment was performed at https://www.genome.jp/kegg/.

### RNA Analysis

Total RNA was isolated with Trizol Reagent (Invitrogen, United States). The reverse transcription of miRNA was performed with miRcute Plus miRNA First-Strand cDNA Kit (Tiangen, Beijing, China) and mRNA reverse transcription was performed with the PrimeScript RT Reagent Kit with gDNA Eraser (Takara, Japan). The qPCR was performed using SYBR Green qPCR Master Mix (Takara, Japan). The qPCR results were analyzed by 2^-ΔΔCt^ method. The gene expression was normalized to the expression of *β-actin*. The miRNA expression was normalized to the expression of *U6*. The primers for qPCR are listed in Table 1.

**Table 1.**
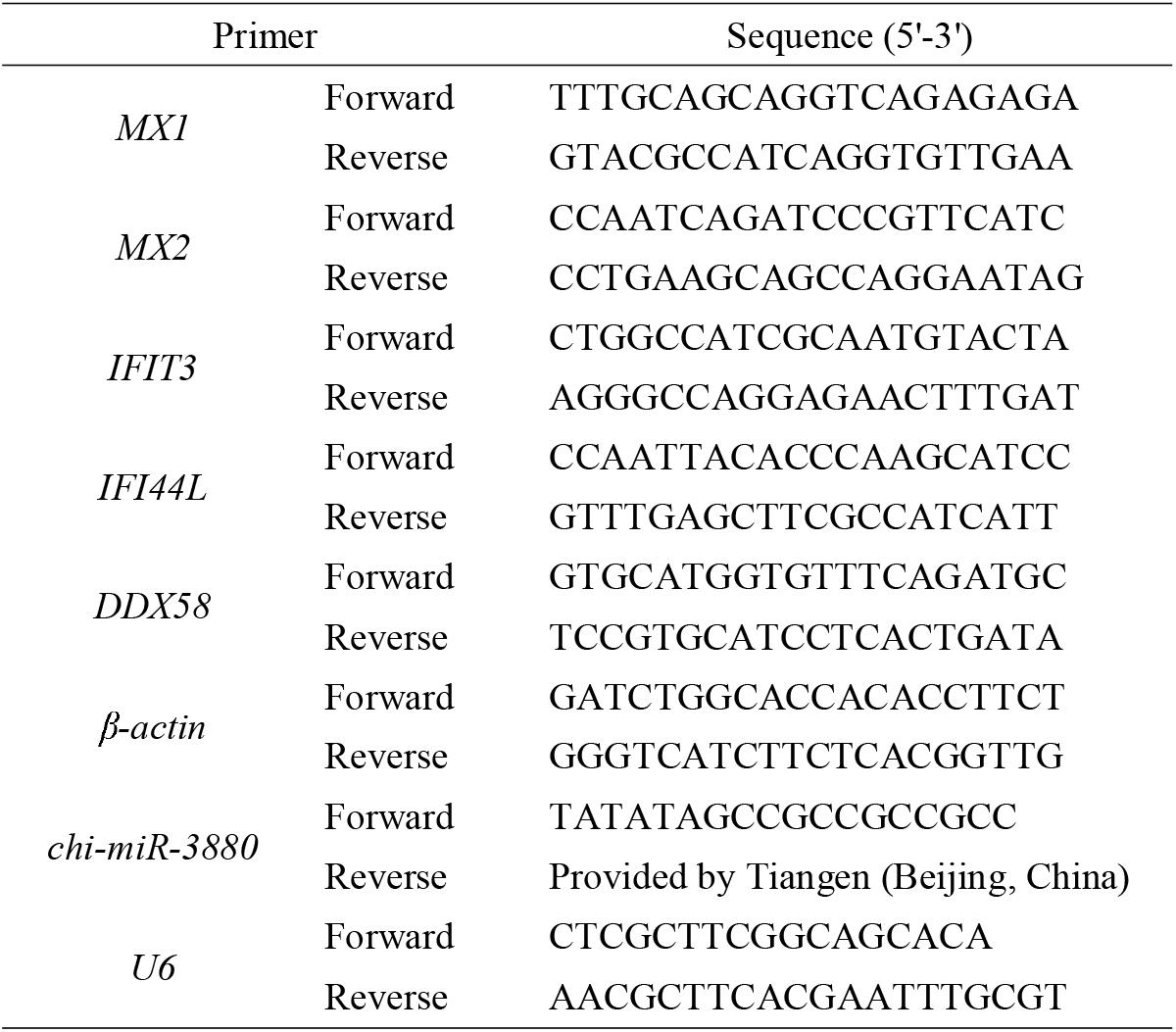
Primers for qPCR

### Flow Cytometry

The cells were gently collected by trypsinization free of EDTA. Annexin V-FITC/PI apoptosis kit (7Sea Biotech, Shanghai, China) was applied to stain cells. Specifically, harvested cell pellet was washed by phosphate buffer saline, and then suspended with 300 μl of 1 X Binding Buffer. The suspension was incubated with 5 μl of Annexin V-FITC in the dark at room temperature for 15 minutes. Then 10 μl of PI was added and incubated with the suspension on ice for five minutes. The samples went through flow cytometer (BD, USA) immediately after incubation. In the results from flow cytometry, upright quadrant represents late apoptosis; downright represents early apoptosis; upleft represents necrosis; downleft represents normal cells.

### Immunohistochemistry

The paraffin-embedded mammary tissues come from the mice previously injected by normal saline or chi-miR-3880 agomir. To be specific, the injection of 10 nmol chi-miR-3880 to postpartum mice was performed every three or four days. The mice were injected six times in total and the tissues were collected at day 22 [7]. The slides with tissue sections were incubated at 68 °C dry oven for 2 hours and deparaffinized in xylene for 20 minutes. The slides went through a gradient of ethanol (100%, 100%, 95%, 80% and 0%) for rehydration; citrate buffer solution for antigen retrieval; 0.5% Triton X-100 for permeation; and 3% hydrogen peroxide to remove endogenous peroxidase. The sections were blocked by goat serum for 20 minutes, and then went to antibody incubation. DAB Substrate kit (Solarbio, DA1010, Beijing, China) was applied as color developer. Haematoxylin was counterstained to show nucleus.

### 5-Ethynyl-20-Deoxyuridine Assay

The cell proliferation was monitored 24 hours post-transfection. Proliferated GMECs in two hours were labeled by BeyoClick 5-Ethynyl-20-Deoxyuridine (EdU) fluorescence in EdU Cell Proliferation Kit with Alexa Fluor 488 (Beyotime, Shanghai, China). The cell nucleus was labeled by DAPI (Beyotime, Shanghai, China). Image-Pro Plus 6.0 Software was used to analyze relative proliferation cells.

### Western Blot

The protein of GMECs was isolated by RIPA lysis buffer with protease inhibitor and phosphatase inhibitor (Thermo Fisher Scientific, USA). The protein was quantified by a BCA protein assay kit (Solarbio, Beijing, China). Equal amount of protein was mixed with SDS-PAGE loading buffer (Solarbio, Beijing, China) and cooked a 98 °C for 10 minutes. The protein was loaded to a SDS-PAGE gel to detect specific protein expression. Primary antibodies applied include: Bax (BBI, D220073, Shanghai, China), Bcl-2 (BBI, D260117, Shanghai, China), p-PI3K p110 beta (Bioss, bs-6417R, Beijing, China), PI3K p110 beta (Bioss, bs-6423R, Beijing, China), AKT (Cell Signaling Technology, 4685, USA), p-AKT (Cell Signaling Technology, 4060, USA), mTOR (Boster Biological Technology, BM4182, USA), p-mTOR (Boster Biological Technology, BM4840, USA), Caspase-3 (Cell signaling technology, 9662, USA), and β-actin (Beyotime, AA128, Shanghai, China). Horse Radish Peroxidase-conjugated IgG secondary antibodies were purchased from Beyotime (Shanghai, China). The immunoblots were exposed and quantified through the Quantity One program (Bio-Rad, California, USA).

### Oil O Staining

The GMECs were fixed with 4% paraformaldehyde for 30 minutes followed by a phosphate buffer saline wash. The cells were incubated in 60% isopropanol for 30 minutes and then exposed to oil O stain solution (Solarbio, Beijing, China). The stain solution were washed off by 60% isopropanol and the cells were further washed by phosphate buffer saline. The density of lipid drops was analyzed by Image-Pro Plus 6.0 Software.

### Statistics

The data were shown as mean ± standard error. All the treated groups were compared to the control group. The experiments were independently repeated at least three times. The significance between groups was analyzed by student t-test. * represents p < 0.05, and ** represents p < 0.01.

## Results

### The analysis of differentially expressed genes (DEGs) induced by chi-miR-3880 in GMEC

To investigate the role of chi-miR-3880, the GO terms and KEGG pathways of downstream DEGs induced by chi-miR-3880 in GMEC were enriched. It was found that the DEGs were involved in 23 GO terms (8 GOs in biological process, 10 GOs in cellular component, and 5 GOs in molecular function). The 23 GO terms are shown as a histogram in Figure 1A, including the immune system process (GO: 0002376) and response to stimulus (GO: 0050896) in biological process. The KEGG enrichment analysis revealed that the DEGs were involved in 26 KEGG pathways (Supplementary File 1). The top 20 KEGG pathways are shown in Figure 1B, including RIG-I-like receptor signaling pathways (ko04622).

**Figure 1.**
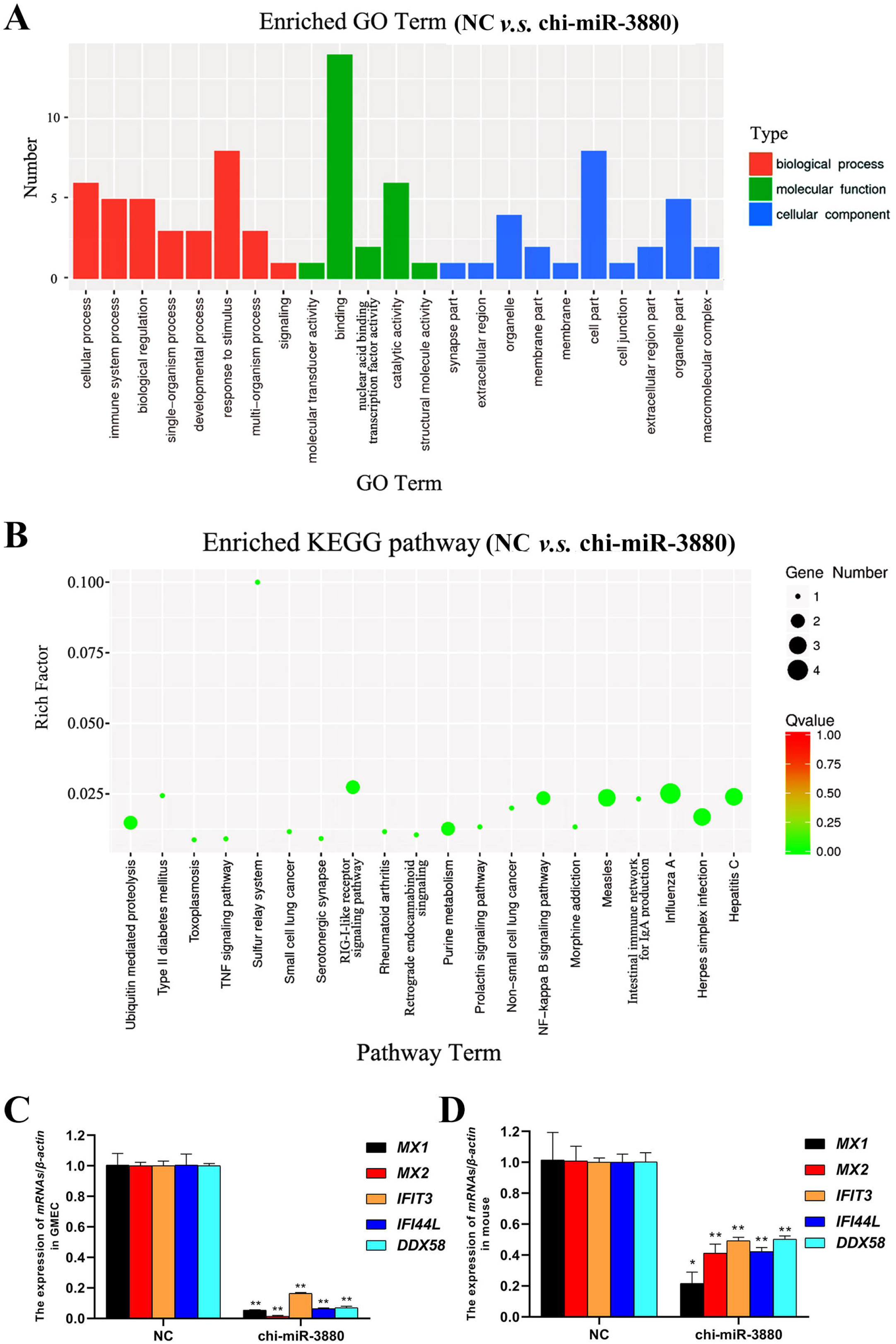
The analysis of differentially expressed genes (DEGs) regulated by chi-miR-3880. (A, B) The GO analysis (A) and KEGG pathway analysis (B) of DEGs regulated by chi-miR-3880; (C, D) The validation of the regulation of interferon stimulated genes (ISGs) by chi-miR-3880 in GMEC (C) and mouse (D).

The expression of the interferon stimulated genes (*MX1, MX2, IFIT3, IFI44L*, and *DDX58*) among the DEGs were detected by RT-qPCR in GMECs transfected by chi-miR-3880 or negative control (NC) to validate the accuracy of RNA sequencing. It shows in Figure 1C that *MX1, MX2, IFIT3, IFI44L*, and *DDX58* were downregulated by chi-miR-3880, which was consistent with the result of RNA sequencing. To investigate the regulation of chi-miR-3880 to the expression of *MX1, MX2, IFIT3, IFI44L*, and *DDX58 in vivo*, the total RNA of mammary gland from mice that had been injected from caudal vein with chi-miR-3880 agomir or normal saline [7] was extracted for RT-qPCR. It showed that the expression of *MX1, MX2, IFIT3, IFI44L*, and *DDX58* were reduced by chi-miR-3880 agomir in mice (Figure 1D).

### The relationship among chi-miR-3880, IFNγ, and *DDX58*

To investigate the regulation between chi-miR-3880 and IFNγ, the expression of chi-miR-3880 and its downstream interferon stimulated genes (*MX1, MX2, IFIT3, IFI44L*, and *DDX58*) was detected in IFNγ (0, 1, 5, 10 ng/ml) treated GMECs. It is shown that the expression of chi-miR-3880 was reduced by IFNγ in a concentration dependent manner (Figure 2A), while the expression of the downstream genes of chi-miR-3880 (*MX1, MX2, IFIT3, IFI44L*, and *DDX58*) was elevated by IFNγ (Figure 2B). *DDX58* was knocked down by siRNA *DDX58* (si*DDX58*) to find out the regulation of *MX1, MX2, IFIT3* and *IFI44L* by *DDX58*. It is shown in Figure 2C that si*DDX58* reduced the expression of *MX1, MX2, IFIT3* and *IFI44L*. The expression of chi-miR-3880 was also slightly decreased by si*DDX58* (Figure 2D). The protein expression of DDX58 regulated by chi-miR-3880 and IFNγ was examined. It showed that chi-miR-3880 decreased the expression of DDX58 (Figure 2E, 2G), while 5 and 10 ng/ml IFNγ increased the expression of DDX58 (Figure 2F, 2G). The DDX58 immunostaining of mammary gland from mice showed that chi-miR-3880 agomir reduced the expression of DDX58 (Figure 2H).

**Figure 2.**
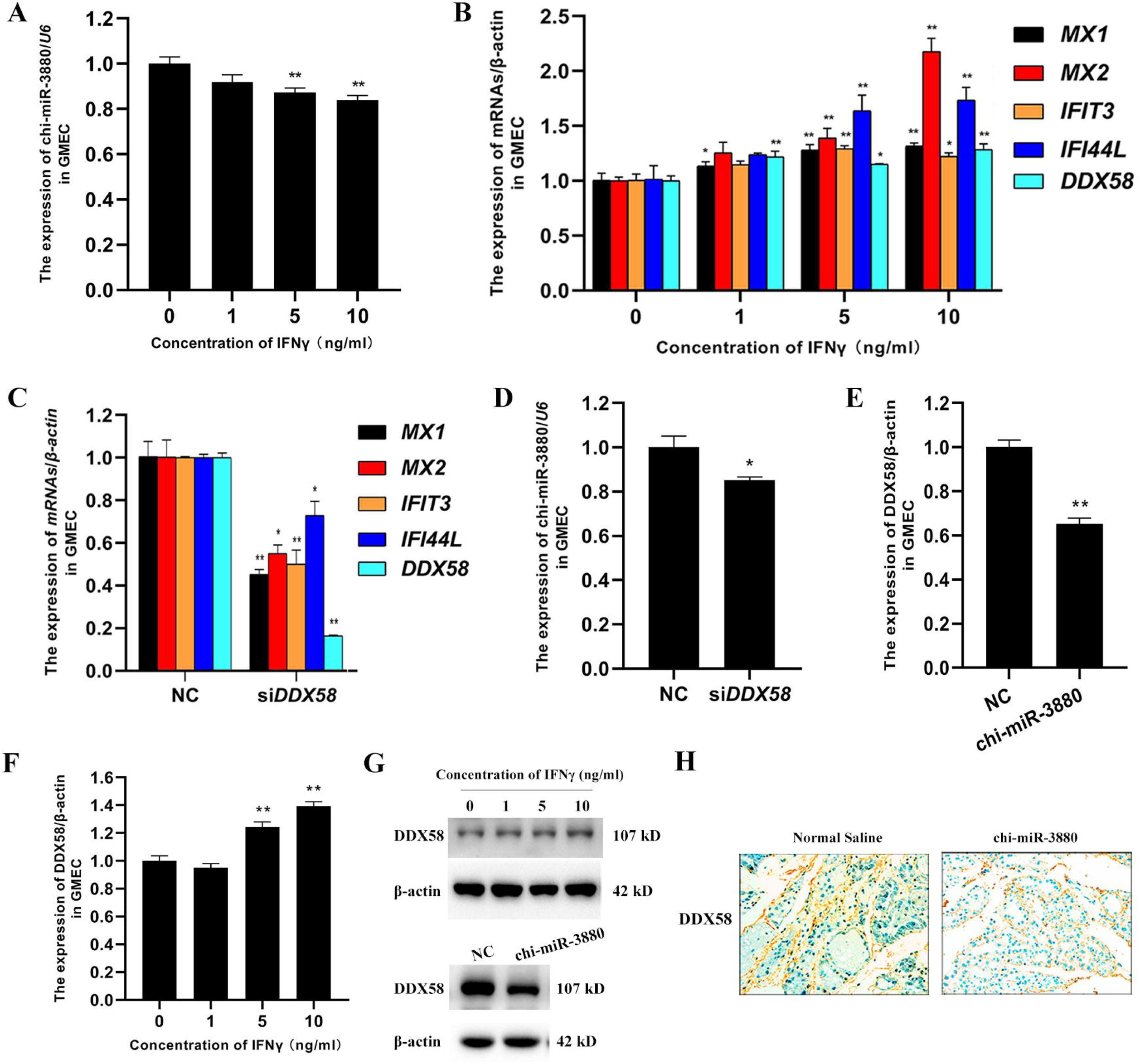
The regulation among IFNγ, chi-miR-3880, and ISGs. (A) IFNγ reduced the expression of chi-miR-3880 in a concentration-dependent manner in GMEC; (B) IFNγ upregulated the expression of the ISGs (*MX1, MX2, IFIT3, IFI44L*, and *DDX58*) in GMEC; (C) The knockdown of *DDX58* resulted in the decreased expression of *MX1, MX2, IFIT3*, and *IFI44L* in GMEC; (D) The knockdown of *DDX58* lowered the expression of chi-miR-3880 in GMEC; (E, G) Chi-miR-3880 decreased the protein expression of DDX58 in GMEC; (F, G) IFNγ lifted the protein expression of DDX58 in a concentration-dependent manner in GMEC; (H) Chi-miR-3880 depressed the expression of DDX58 in mouse mammary gland.

### IFNγ/chi-miR-3880/*DDX58* axis regulates GMEC proliferation and apoptosis

The regulation of GMEC proliferation and apoptosis by IFNγ and *DDX58* was then investigated. When exposed to IFNγ, the proliferation of GMEC was inhibited (Figure 3A, 3B), and the late apoptotic rate of GMEC was increased (Figure 3E, 3F). It was shown that si*DDX58* promoted GMEC proliferation (Figure 3C, 3D) and suppressed GMEC late apoptosis (Figure 3G, 3H). The expression of apoptosis-related protein, such as cleaved-Caspase3/pro-Caspase3 and Bcl-2/Bax, was detected in GMEC to explore through which pathways IFNγ/chi-miR-3880/*DDX58* axis regulated apoptosis. The immunoblots revealed that IFNγ triggered the cleavage of Caspase3 and decreased the expression of Bcl-2 (Figure 3I, 3J), while si*DDX58* reduced the expression of cleavaged-Caspase3 and increased the expression of Bcl-2 (Figure 3K, 3L). It is shown in Figure 3M, 3N that chi-miR-3880 decreased the expression of cleavaged-Caspase3.

**Figure 3.**
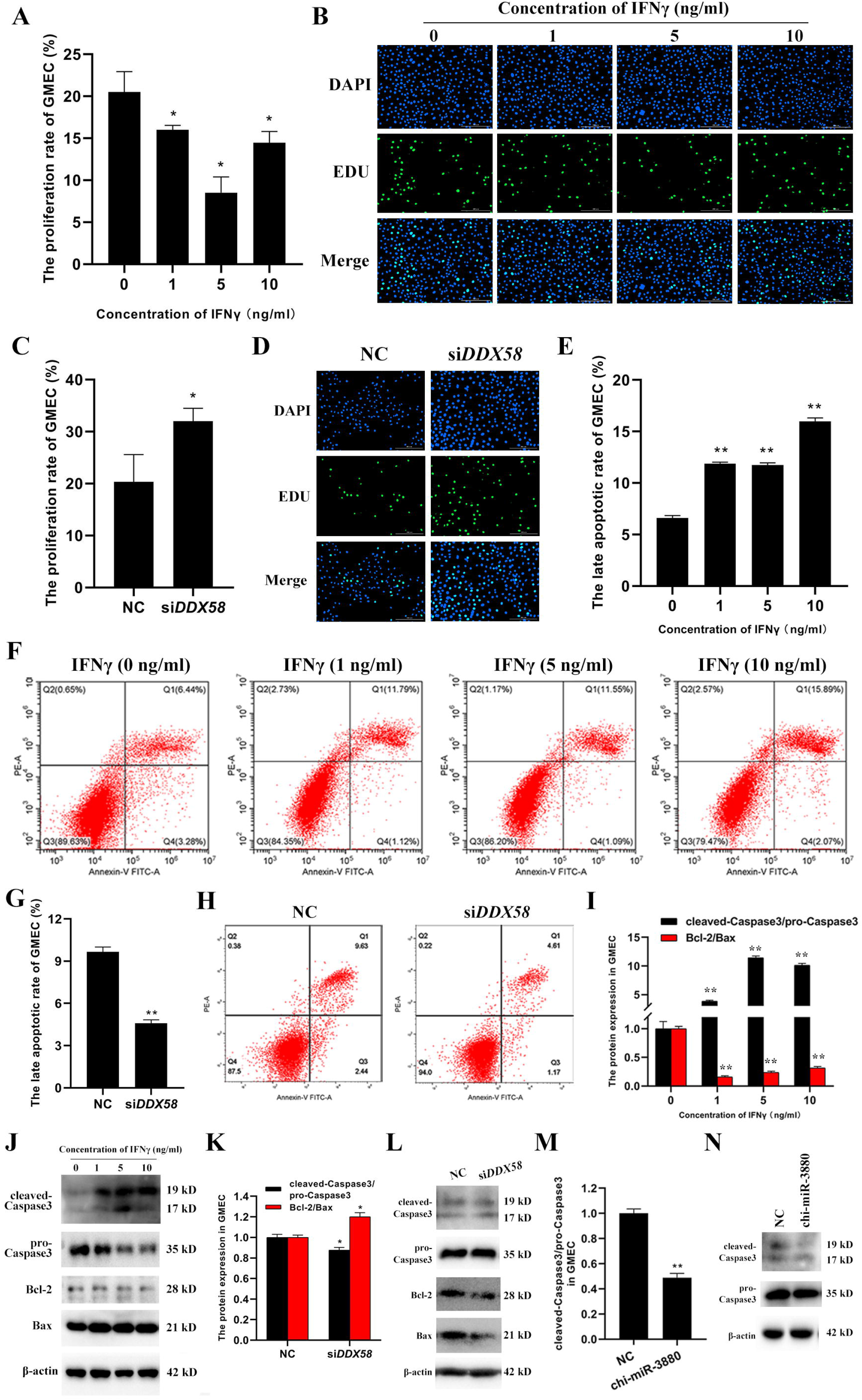
The regulation of proliferation and apoptosis of GMEC by IFNγ/chi-miR-3880/*DDX58* axis. (A-D) IFNγ suppressed (A, B) while si*DDX58* promoted (C, D) the proliferation of GMEC; (E-H) IFNγ facilitated (E, F) while si*DDX58* inhibited (G, H) the apoptosis of GMEC; (I, J) IFNγ drove the activation of Caspase-3 and restrained the expression of Bcl-2/Bax; (K, L) The knockdown of *DDX58* decreases the activation of Caspase-3 and facilitated the expression of Bcl-2/Bax; (M, N) Chi-miR-3880 repressed the activation of Caspase-3.

### IFNγ/chi-miR-3880/*DDX58* axis contributes to the regulation of mammary involution and lipid accumulation

To explore the role of IFNγ and *DDX58*, the lipid accumulation in GMEC was examined by oil O staining. It is shown in Figure 5 that IFNγ impaired the lipid accumulation (Figure 4A, 4C), and si*DDX58* enhanced the lipid accumulation (Figure 4B, 4C). To find out through which pathway IFNγ and *DDX58* regulated lipid accumulation, the expression of phosphorylated PI3K, AKT and mTOR was detected. As shown in Figure 4D–4F, the phosphorylation of PI3K, AKT and mTOR was decreased by IFNγ treatment, and increased by si*DDX58*.

**Figure 4.**
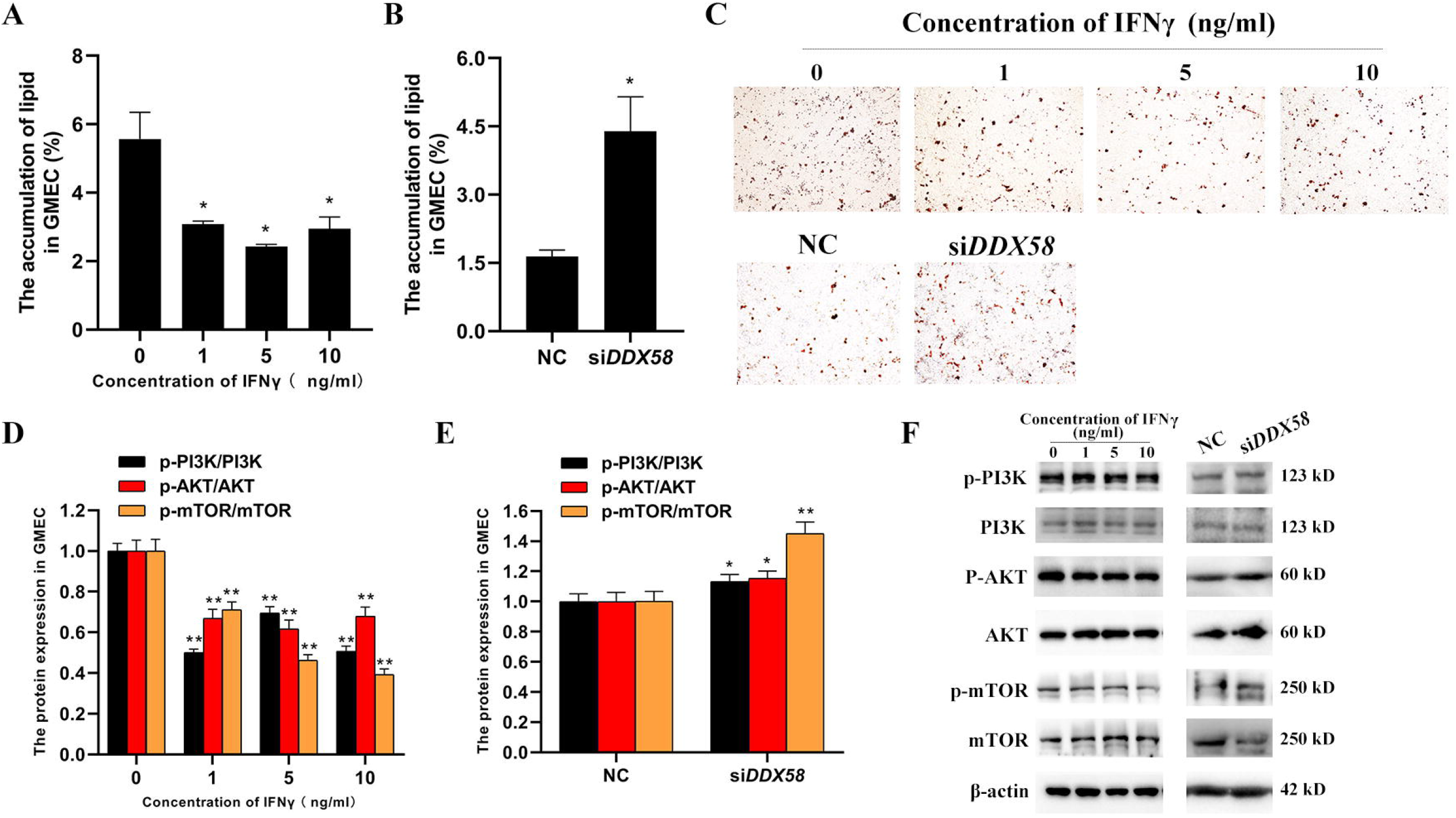
IFNγ and *DDX58* modulate lipid accumulation and the activation of PI3K/AKT/mTOR pathway. (A-C) IFNγ suppressed (A, C) while si*DDX58* promoted (B, C) the lipid accumulation in GMEC; (D-F) IFNγ restrained (D, F) while si*DDX58* contributed to (E, F) the phosphorylation of PI3K/AKT/mTOR pathway.

**Figure 5.**
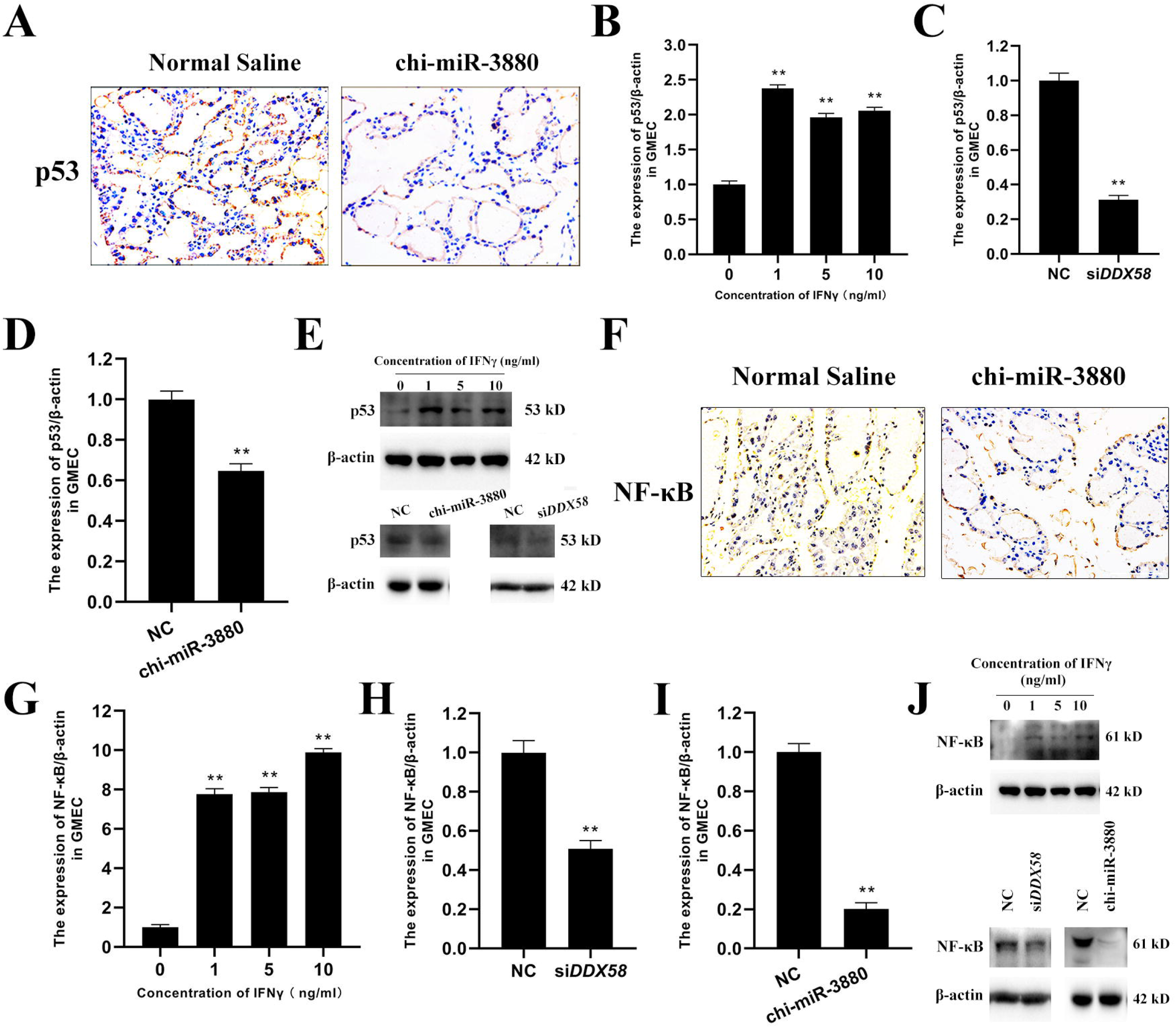
The modulation of the expression of involution markers by IFNγ/chi-miR-3880/*DDX58* axis. (A) Chi-miR-3880 inhibited the expression of p53 in the mammary gland of mouse; (B-E) IFNγ increased (B, E) while si*DDX58* (C, E) and chi-miR-3880 (D, E) decreased the expression of p53 in GMEC; (F) Chi-miR-3880 inhibited the expression of NF-κB in the mammary gland of mouse; (G-J) IFNγ lifted (G, J) while si*DDX58* (H, J) and chi-miR-3880 (I, J) reduced the expression of NF-κB in GMEC.

To evaluate the effect of chi-miR-3880 on mammary status, p53 and NF-κB expressions were detected. It was found that the expression of p53 and NF-κB was decreased in mammary gland of mice in chi-miR-3880 agomir group compared to normal saline group (Figure 5A, 5F). In GMEC, the protein expression of p53 and NF-κB was upregulated by IFNγ and downregulated by chi-miR-3880 and si*DDX58* (Figure 5B–5E, 5G–5J).

## Discussion

In this study, the relationship among chi-miR-3880, IFNγ, and *DDX58*, as well as their role in mammary gland were investigated. During peak lactation period of Guanzhong dairy goats, the expression of chi-miR-3880 goes over two times higher than colostrum period [15]. In our previous study, it is revealed that chi-miR-3880 contributes to the proliferation and lipid accumulation of goat mammary epithelial cells (GMECs) and to the development of the mammary gland during lactation [7]. Interestingly, RNA-seq analysis revealed that quite a few interferon-stimulated genes (ISGs) are downregulated by chi-miR-3880, such as *MX1, MX2, IFIT3, IFI44L*, and *DDX58* [7]. The downregulation of *MX1, MX2, IFIT3, IFI44L*, and *DDX58* by chi-miR-3880 was validated by qPCR in this study. To investigate the regulation of chi-miR-3880 in GMEC, the downstream genes of chi-miR-3880 were enriched to GO terms and KEGG pathways. It was shown that the downstream genes were enriched in GO terms of immune system process and response to stimulus and RIG-I-like receptor signaling pathway. Interferon is involved in the priming of immune cells and the training of innate immune memory, and also affect ISGs [13]. Recently, IFNγ is identified as a potential marker of caseous lymphadenitis in lactating sheep [14]. The reduced expression of chi-miR-3880 regulated by IFNγ (Figure 2A) indicates that chi-miR-3880 might participate in the immunity and inflammation. Strikingly, IFNγ enhanced the expression of ISGs downregulated by chi-miR-3880, indicating an axis of IFNγ, chi-miR-3880 and ISGs. Among the ISGs, the knockdown of *DDX58* led to the downregulation of the other four ISGs (*MX1, MX2, IFIT3, IFI44L*), which suggests the major role of *DDX58* in the detected ISGs. The proliferation inhibition and apoptosis promotion in GMEC led by IFNγ and *DDX58* demonstrates that the regulation of proliferation and apoptosis by chi-miR-3880 could be realized in IFNγ/chi-miR-3880/*DDX58* axis through Caspase3 activation and Bcl-2/Bax pathway. The lipid accumulation in GMECs was modulated by IFNγ/chi-miR-3880/*DDX58* axis as well through the activation of PI3K/AKT/mTOR pathway. It was found that chi-miR-3880 reduced the expression of NF-κB and p53 both in vitro and in vivo. The involvement of IFNγ and *DDX58* in the regulation of NF-κB and p53 was also confirmed in GMEC. As upregulated NF-κB and p53 are identified as signs of involution process in lactation [16, 17], it is likely that IFNγ/chi-miR-3880/*DDX58* axis could modify involution. Overall, the study shows that the modulation of chi-miR-3880 in mammary involution, proliferation, apoptosis and lipid accumulation of GMEC could be achieved in IFNγ/chi-miR-3880/*DDX58* axis (summarized in Figure 6).

**Figure 6.**
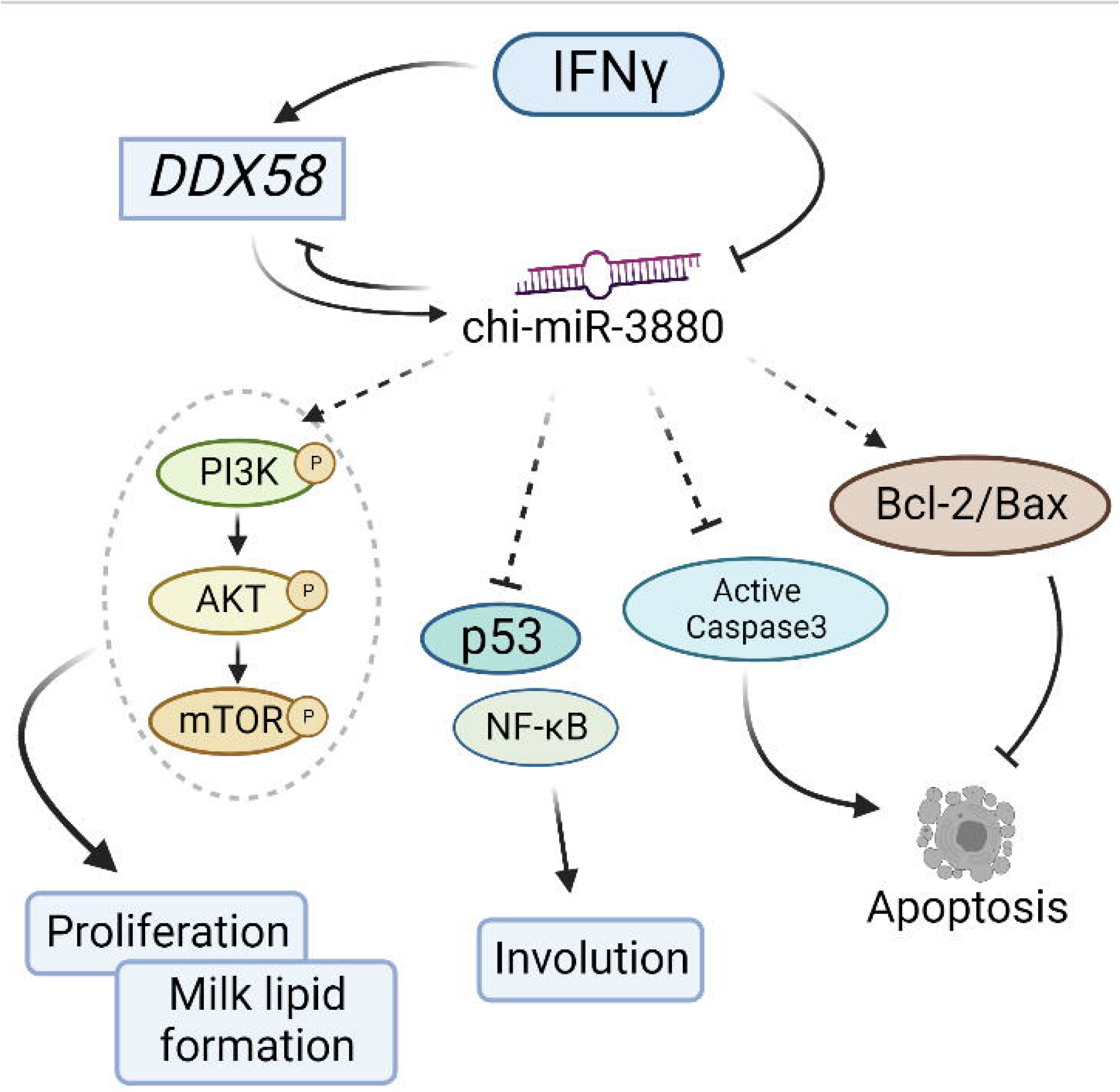
A sketch showing the idea of the study.

IFNγ is an inflammatory cytokine that alters immune-related miRNAs [18, 19], indicating the importance of studying the interaction between IFNγ and miRNAs. High IFNγ production contributes to the activation of immune system during dry lactation stage, and is related to the onset of diseases postpartum [20]. The study on IFNγ-miRNA regulatory pathway would be helpful to understand IFNγ-related diseases.

## Acknowledgement

Thanks for the supports from the National Natural Science Foundation of China (81860762), Scientific Research Guiding Plan Topic of Qinghai Hygiene Department (2018-wjzdx-131), Shaanxi Science and Technology Innovation Project Plan (2017ZDXM-NY-081 and 2018ZDCXL-NY-01-04), Shaanxi Key Research and Development Program (2020ZDLNY02-01 and 2020ZDLNY02-02), and Natural Science Foundation of Shaanxi Province (2020JQ-868). Thanks for the help of Biorender (https://app.biorender.com/) in drawing Figure 6. Thanks for the help of Servicebio Technology Company (Wuhan, China) on immunohistochemistry.

